# Conformational basis of subtype-specific allosteric control of NMDA receptor gating

**DOI:** 10.1101/2024.02.10.579740

**Authors:** Julia Bleier, Philipe Ribeiro Furtado de Mendonca, Chris Habrian, Cherise Stanley, Vojtech Vyklicky, Ehud Y. Isacoff

## Abstract

N-methyl-D-aspartate receptors are ionotropic glutamate receptors that are integral to synaptic transmission and plasticity. Variable GluN2 subunits in diheterotetrameric receptors with identical GluN1 subunits set very different functional properties, which support their individual physiological roles in the nervous system. To understand the conformational basis of this diversity, we assessed the conformation of the common GluN1 subunit in receptors with different GluN2 subunits using single-molecule fluorescence resonance energy transfer (smFRET). We established smFRET sensors in the ligand binding domain and modulatory amino-terminal domain to study an apo-like state and partially liganded activation intermediates, which have been elusive to structural analysis. Our results demonstrate a strong, subtype-specific influence of apo and glutamate-bound GluN2 subunits on GluN1 rearrangements, suggesting a conformational basis for the highly divergent levels of receptor activity, desensitization and agonist potency. Chimeric analysis reveals structural determinants that contribute to the subtype differences. Our study provides a framework for understanding GluN2-dependent functional properties and could open new avenues for subtype-specific modulation.

## Introduction

Fast glutamatergic neurotransmission mediated by ionotropic glutamate receptors forms the basis of excitatory neurotransmission in the mammalian central nervous system. The ionotropic glutamate receptor family includes several subtypes including α-amino-3-hydroxy-5-methyl-4- isoxazolepropionic acid (AMPA), kainate, and N-methyl-D-aspartate (NMDA) receptors, which not only detect release of glutamate into the synaptic cleft, but also act as sensitive filters of the signal received; each type responds to different ranges of agonist concentration, differentially depends on co-agonists, modulators and voltage, and generates membrane currents with distinct cation permeabilities, rise and fall kinetics and propensities to desensitize.^1^ Whereas AMPA and kainate receptors primarily mediate fast synaptic events, opening quickly and desensitizing quickly and completely, NMDA receptors, have higher affinity for glutamate, require glycine or D-serine as a co-agonist, desensitize less, and deactivate more slowly.^2^ NMDA receptors contain two pairs of alternating obligate GluN1 and variable GluN2 or GluN3 subunits, each consisting of an intracellular C-terminal domain (CTD), a pore-forming transmembrane domain (TMD), and a large extracellular domain made up of a ligand-binding domain (LBD) and an N-terminal domain (NTD).^3,4^ Glutamate binding to the GluN2 LBD and glycine or D-serine binding to the GluN1 LBD are loosely coupled to channel opening,^5–7^ but even in saturating concentrations of agonists, the channel pore spends much of the time closed.^8–10^

NMDA receptors with different GluN2 subunits exhibit distinct spatiotemporal expression patterns in the developing brain, as well as distinct subcellular localization, roles in synaptic transmission, plasticity, and disease, and they differ in several loosely covariant physiological properties.^2,11–14^ GluN2A NMDAR have the highest single channel open probability (P_o_; ∼50%), followed by GluN2B and GluN2C, with GluN2D having the lowest P_o_ (∼1%).^2^ Subtypes with lower P_o_ exhibit less desensitization and slower deactivation,^15–18^ and higher potency for both glutamate^19–22^ and glycine.^23,24^ This provides a physiological framework for transmitting diverse synaptic signals: low concentrations of glutamate, as seen in spillover from neighboring synapses, can activate low P_o_ receptors with slow kinetics and high-concentration glutamate transients in the synaptic cleft can activate higher P_o_ receptors briefly in a precisely timed manner. Single channel patch-clamp recordings have been used to construct kinetic schemes for pathways between multiple non-conducting closed and cation-permeable open states,^8,25–29^ though only transitions that open and close the pore are observed. Electrophysiological study of receptors with chimeric GluN2 subunits has proven successful in identifying structural regions that contribute to observed properties.^24,30–32,22^ Structural studies have transformed our understanding of NMDAR conformation by capturing each diheterotetrameric subtype at high resolution.^3,4,33–39^ Though distinct conformations have been observed in different subtypes and in different ligand conditions, it remains difficult to understand how subtype-specific gating emerges without observing the conformations adopted in each subtype in the resting state and with glycine or glutamate alone.

To elucidate the basis of functional diversity between NMDA receptor subtypes, we sought to study the conformational pathway of NMDA receptor activation, from the resting state and through rearrangements elicited by each co-agonist individually. To do this, we turned to single-molecule fluorescence energy transfer (smFRET), which has been used to capture conformational pathways in ion channels and neurotransmitter receptors, including NMDA receptors.^40–45^ We employed smFRET to measure distances between identical sites in the two GluN1 subunits— either in the NTD or LBD— in the resting state and in subunit-specific orthosteric agonists. We compared structural rearrangements of the same GluN1 in receptors containing unlabeled versions of either GluN2A, B, C or D. Our experiments revealed dramatic GluN2 subtype differences between the conformational dynamics of the GluN1 NTD and LBD in resting and partially liganded states and a key difference in the GluN1 NTD in the fully liganded state. Chimeric analysis identified extracellular domain components responsible for these subtype differences. Our observations provide a conformational basis for the physiological diversity among NMDA receptor subtypes.

## Results

In order to assess ligand-gated conformational rearrangements of the extracellular domain of NMDA receptors, we monitored the proximity of the two GluN1 subunits under multiple ligand conditions. To understand how these conformational rearrangements vary between subtypes with different functional properties, we combined identical, labeled GluN1 subunits with unlabeled versions of each of the GluN2 subtypes (GluN2A, GluN2B, GluN2C or GluN2D). We used unnatural amino acid (UAA) incorporation in the GluN1 subunit to introduce a chemical handle that could be site-specifically labeled with FRET donor and acceptor fluorophores (**Fig. S1A**). GluN2 subunits with a C-terminal Human influenza hemagglutinin (HA) tag were co-expressed in HEK293T cells with a GluN1-1a subunit containing an amber stop codon at the desired UAA incorporation site, along with a plasmid containing Pyrrolysyl-tRNA Synthetase(AF) and its cognate *M. mazei* tRNA^Pyl^, which recognizes the amber stop codon.^46^ To increase incorporation efficiency, we modified the previously published construct by inserting a nuclear export signal,^47^ the woodchuck hepatitis virus post-transcriptional regulatory element,^48^ three additional copies of the tRNA and a translation elongation factor. Together these elements should increase mRNA export into the cytoplasm and decrease its degradation, increase tRNA availability and facilitate translational elongation. Incubation of transfected cells with the UAA, trans-Cyclooct-2-en–L-Lysine (TCOK), enabled its incorporation in the position of the amber stop codon. Subsequent stochastic labeling with pyrimidyl-tetrazine-conjugated Alexa Fluor-555 (FRET donor) and Alexa Fluor-647 (FRET acceptor) dyes was achieved through bioorthogonal click chemistry. We verified that labeled receptors trafficked to the plasma membrane and retained their ion channel function (**Fig. S1D,E**).

To understand conformational changes across the extracellular domain we selected UAA incorporation sites in the R1 lobe of the NTD, GluN1-1a(W56TAG), and the D2 lobe of the LBD, GluN1-1a(D677TAG) (**Fig. S1C**). Transfected HEK293T cells were labeled, lysed and receptors containing the HA-tagged GluN2 subunit were immune-purified in the one-step SiMPull method^42,49^ by immune-trapping onto a PEG-passivated coverslip containing a sparse lawn of PEG-biotin to which neutravidin and biotinylated anti-HA antibody were bound (**Fig. S1B**). Total internal reflection fluorescence microscopy was used to visualize single fluorescent molecules on individual receptors and to select for analysis receptors with one FRET donor and one acceptor. The FRET efficiency of receptors labeled at these sites indicates the proximity of the two GluN1 NTDs and lower LBDs resulting from large-scale differences in extension of the GluN1/GluN2 dimers as well as rotations of the LBD and NTDs. For each region of interest imaged, FRET traces were combined into FRET efficiency histograms; histograms in figures represent the average of multiple regions of interest (see Methods). Comparing these efficiencies across ligand conditions and receptor subtypes allowed us to understand ligand-gated conformational changes and how they are influenced by each GluN2 subunit.

### Agonist-induced convergence of GluN1 LBD conformation

To study the rearrangement of the GluN1 LBD, we monitored the inter-LBD distance between GluN1 subunits labeled at GluN1-1a(D677TAG) in receptors containing either GluN2A, B, C or D subunits. GluN1-1a residue 677 is located in the D2 (lower) lobe of the LBD and should provide a measure of agonist-induced closure as well as the wide range of rotations that the LBD can undergo (**Fig. S1C**). Despite their distinct physiological properties, structural studies have shown that GluN1/GluN2 receptors, with both glutamate and glycine bound, can adopt globally similar conformations across subtypes.^36–39^ Consistent with this, we find that, in saturating concentration of glutamate (1 mM) and glycine (100 μM), GluN1-1a(D677TAG) populates similar well-defined conformations for GluN2A, B, C and D with median FRET efficiency ∼0.25 (**Fig. 1C**). The similarity in GluN1 LBD FRET distributions between fully liganded receptors with different GluN2s suggests a common closed and rotated GluN1 LBD conformation that is permissive to channel opening. We then assessed GluN2 dependence of GluN1 conformational dynamics in the activation pathway that leads to this permissive state. We began with the resting state without either co- agonist.

**Figure 1.**
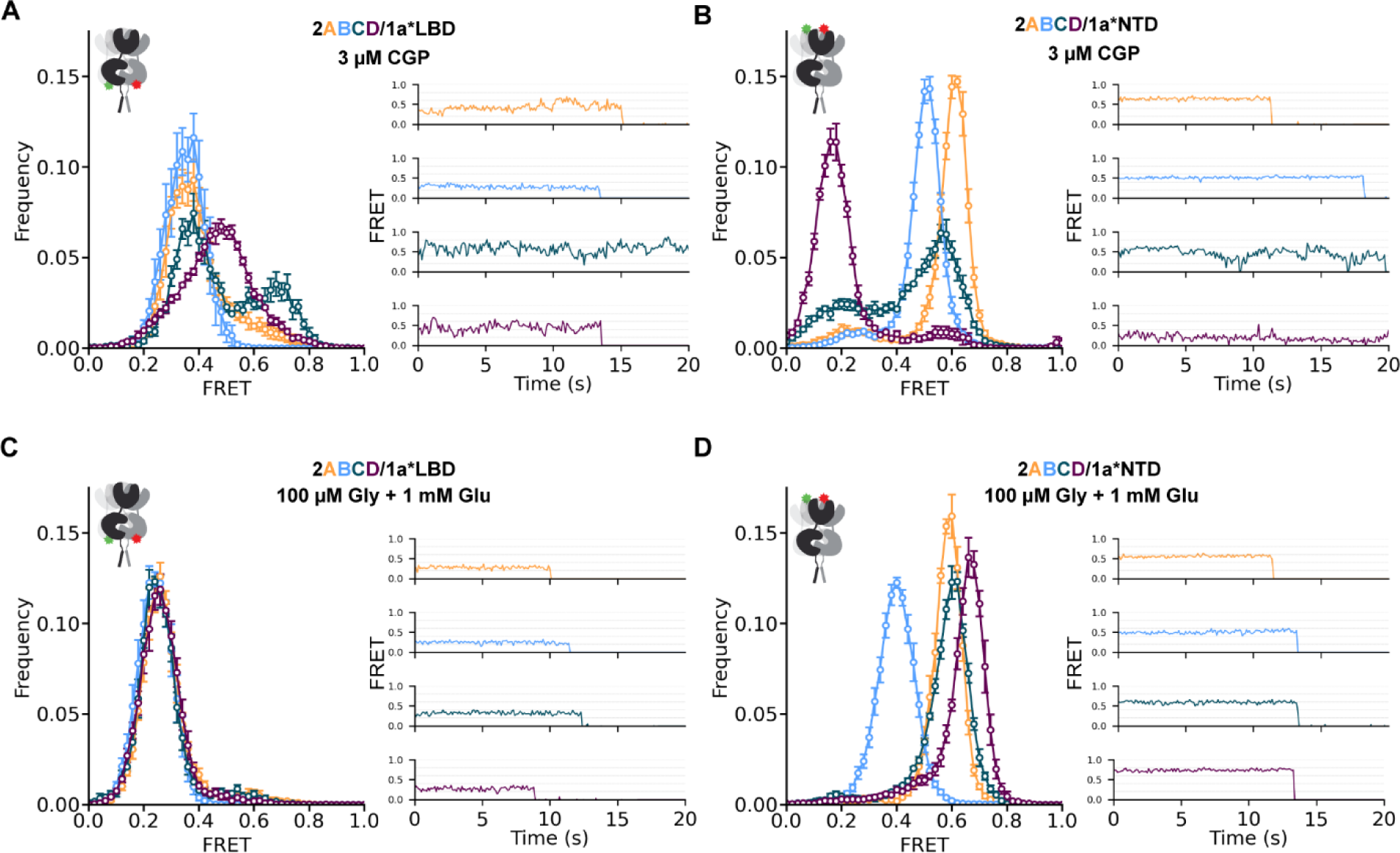
Agonists drive convergence of GluN1 conformation from GluN2 dependent resting states. A-D) Ligand-dependent conformations determined from inter-subunit FRET between GluN1 NTDs with fluorophores incorporated at site W56TAG (A,C) or GluN1 LBDs with fluorophores incorporated at site D677TAG (B,D) in an apo-like state (zero added glycine, 3 μM GluN1 antagonist CGP78608, zero added glutamate) (A,B) or with saturating concentrations of agonists (100 μM Glycine, 1 mM Glutamate) (C,D) when GluN1 is combined with either GluN2A (orange), GluN2B (blue), GluN2C (teal) or GluN2D (magenta). Left) Histograms of smFRET distributions indicating mean and S.E.M. Right) Example smFRET traces with colors corresponding to histogram keys. Labeling sites indicated with green and red stars on cartoons. Total receptor number for each condition listed in Table S1. Donor (Alexa Fluor 555) and acceptor (Alexa Fluor 647) dyes imaged at 10 fps.

The apo conformation of NMDA receptors has been elusive in structural studies and is inherently undetectable in electrophysiological approaches. Regardless of approach, due to the high affinity binding to GluN1, low levels of contaminating glycine must be surmounted to assay an apo-like state. We previously showed that the glycine-site competitive antagonist, CGP78608 (CGP), in the absence of any GluN2 ligand stabilizes an apo-like resting conformation in GluN2B receptors.^45^ Here we used CGP to assess the resting conformation of the GluN1 LBD in the context of each GluN2 subunit. In the GluN1 antagonist-bound / GluN2 empty resting state (3 μM CGP), we observe broader FRET distributions shifted to higher values of FRET efficiency than seen in the fully liganded (1 mM Glu + 100 μM Gly) state for all of the GluN2 subtypes (**compare Fig. 1A to C**). Moreover, the FRET distributions show that the inter-subunit distance between GluN1 lower lobes depends on the identity of the unliganded GluN2 subunit. Gaussian fits to the FRET distributions enable us to estimate relative occupancies among these NMDA receptor subtypes of inferred low (0.35), medium (0.50) and high (0.70) FRET states in the resting state (**Fig. S3**). Occupancy of the low FRET (.35) conformation, which likely corresponds to a pre-activated state with greatest tension exerted on the S2-M3 linker to the channel gate, followed the sequence GluN2B > GluN2A > GluN2C > GluN2D. This suggests that the unliganded GluN2 subunit differentially inhibits GluN1 LBD rotation to the active pre-open conformation, with the strongest inhibition by GluN2D, the receptor with the lowest P_o_.

### GluN2-dependent GluN1 NTD resting splaying and liganded compaction

Structural studies have demonstrated that, primarily under conditions of inhibition,^34,38,50^ NMDA receptors can adopt splayed conformations, in which lower LBDs are closely apposed, upper LBD interfaces are ruptured, and NTDs are moved far apart. We therefore hypothesized that closer apposition of the lower lobe of the GluN1 LBD (*i.e.* higher FRET lower LBD conformations) might be accompanied by more splayed conformations of the GluN1 NTD. We find that, in the apo-like state with CGP bound to the GluN1 LBD, the proximity of the GluN1 NTDs differs greatly between receptors with the different GluN2 subunits, with the greatest GluN1 NTD-NTD distance in GluN2D (median FRET = 0.18), followed by GluN2C (median FRET = 0.48), GluN2B (median FRET = 0.51), and the most proximal in GluN2A (median FRET = 0.60) (**Fig. 1A**). The NTD-NTD distance roughly mirrors the distance between lower LBDs, ie. greater distance (lower FRET) between NTDs corresponds to shorter distance between (higher FRET due to less rotation of) LBDs (**compare Fig. 1A and B**), consistent with coordinated rearrangements of these domains between splayed and compact conformations. The trend in occupancy of splayed conformations in the resting state (GluN2D > GluN2C > GluN2B > GluN2A) matches trends observed in physiological properties. In the fully liganded (1 mM Glu + 100 μM Gly) state, the NTDs also occupy distinct conformations in receptors with different GluN2s (**Fig. 1D**). Notably, we observe that GluN2D, the subtype with the lowest open probability, is super-compact (higher FRET) compared to GluN2A, the subtype with the highest open probability, consistent with what is observed in structural studies.^38,39^

### GluN2 dependence of single-agonist GluN1 conformation

To understand the conformational activation pathway between resting and fully agonist-bound states, we assessed the individual effects of glycine and glutamate on GluN1 conformation. In all subtypes, compared to CGP (3 μM), glycine alone (100 μM Gly) decreases FRET between lower lobes of the GluN1 LBDs, consistent with LBD closure and rotation (**Fig. 2A-D, blue**). The magnitude of the conformational change induced by glycine is greater in GluN2C and GluN2D receptors than in GluN2A and GluN2B receptors. Agonist binding to the GluN2 subunit alone (1 mM Glu + 3 μM CGP) also drives GluN1 LBD rotation, but to a much smaller degree (**Fig. 2A-D, purple**). The largest glutamate-induced GluN1 LBD rotation is seen in the GluN2D receptor, suggesting a particularly strong influence of GluN2D on the GluN1 LBD.

**Figure 2.**
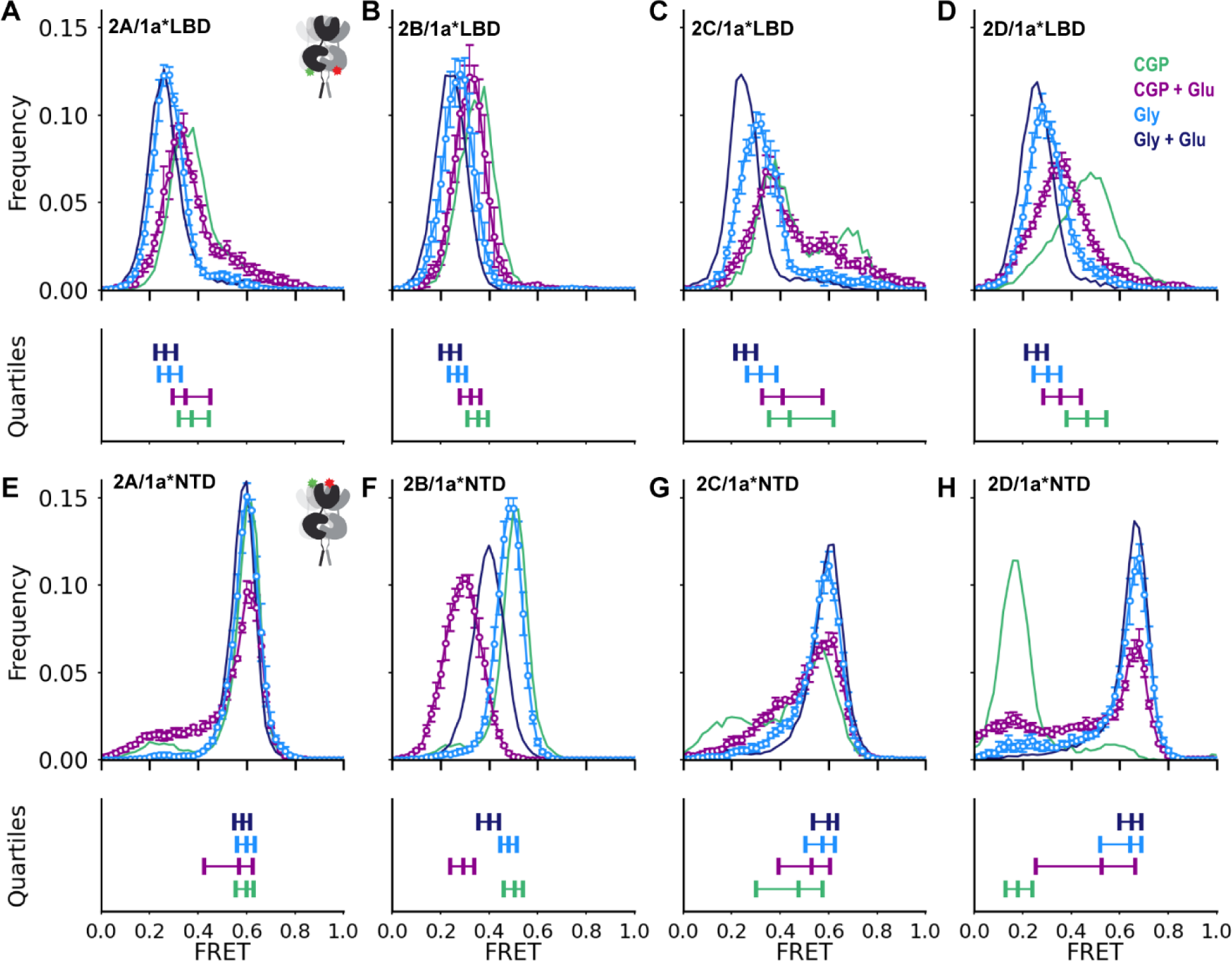
GluN2 dependent regulation of GluN1 conformation in single-agonists. (A-H) Ligand-dependent conformations determined from inter-subunit FRET between GluN1(D677TAG) lower LBD (A-D) or GluN1(W56TAG) NTD (E-H) paired with GluN2A (A,E), GluN2B (B,F), GluN2C (C,G), or GluN2D (D,H) in 3 μM CGP78608 and 1 mM glutamate (purple), 100 μM glycine (blue) in apo-like (zero added glycine, 3 μM GluN1 antagonist CGP, zero added glutamate) (green) and saturating agonist (100 μM Glycine, 1 mM Glutamate) conditions (navy) reproduced from Figure 1. (Upper) Histograms of smFRET distributions, indicating mean and S.E.M. for those not shown in Figure 1. Labeling sites indicated with green and red stars on cartoon insets (A, E). Total receptor number for each condition listed in Table S1. (Lower) Quantification of the spread of the distributions in above histograms, using the same color scheme. For each, the left and right vertical ticks indicate the first and third quartiles and the middle the median. Donor (Alexa Fluor 555) and acceptor (Alexa Fluor 647) dyes imaged at 10 fps. See also Figure S3, 4.

At the level of the NTD, glycine alone, binding to the GluN1 LBD, favors occupancy of the highest FRET, compact, state in each of the receptor subtypes (**Fig. 2E-H, blue**), indicating stabilization of compact states. Glutamate alone, binding to GluN2C or GluN2D, also increases occupancy of compact conformations (**Fig. 2G,H, purple**), with frequent and heterogeneous transitions between splayed, compact, and super-compact conformations observed in GluN2D receptors (**Fig. S4**). However, glutamate binding to GluN2A and GluN2B has the opposite effect, decreasing occupancy of compact conformations (**Fig. 2E,F**). This is consistent with the known apparent negative cooperativity between glutamate and glycine binding in GluN2A receptors,^43,51,52^ and suggests that GluN2B receptors share that negative cooperativity, but that in GluN2C and GluN2D receptors, glutamate instead exerts a positive cooperative effect on GluN1 subunits.

### GluN1-GluN2 coupling depends on the GluN2 NTD α5 helix and GluN1 LBD loop 2

To determine which structural regions in GluN2 subunits are responsible for their distinct allosteric regulation of GluN1, we generated chimeras from GluN2B and GluN2D (**Figs. 3A,S2**), which have the most distinct resting-state conformations and glutamate-induced rearrangements. Swapping the GluN2D NTD into GluN2B [2B(2D NTD)] increased NTD compactness in both the resting (3 μM CGP) and saturating glutamate (1 mM Glu + 3 μM CGP) states so that the glutamate state, but not the resting state, was similar to that of GluN2D (**Fig. 3B)**. To narrow down the critical region for glutamate state regulation, we split the NTD into three regions (N1-3). GluN2B with the GluN2D N1 and N2 or N2 alone did not express well enough to assay. GluN2B with the GluN2D N2 and N3 [2B(2D N2, N3)] showed conformations closest to those seen with transfer of the entire NTD (**Figs. 3B**), suggesting that N2 plays an important role. To test this idea, we substituted individually each of the three alpha helices contained in N2 (α5, α6, α7). Of the three helices, replacement of only the GluN2B α5 helix with that of GluN2D produced the biggest shift to a higher glutamate-alone FRET distribution, and so most closely approached the behavior of the swap of the entire GluN2D NTD (**Fig. 3B**). We also observe that each of the chimeras impacted the compactness of the resting state (3 μM CGP).

**Figure 3.**
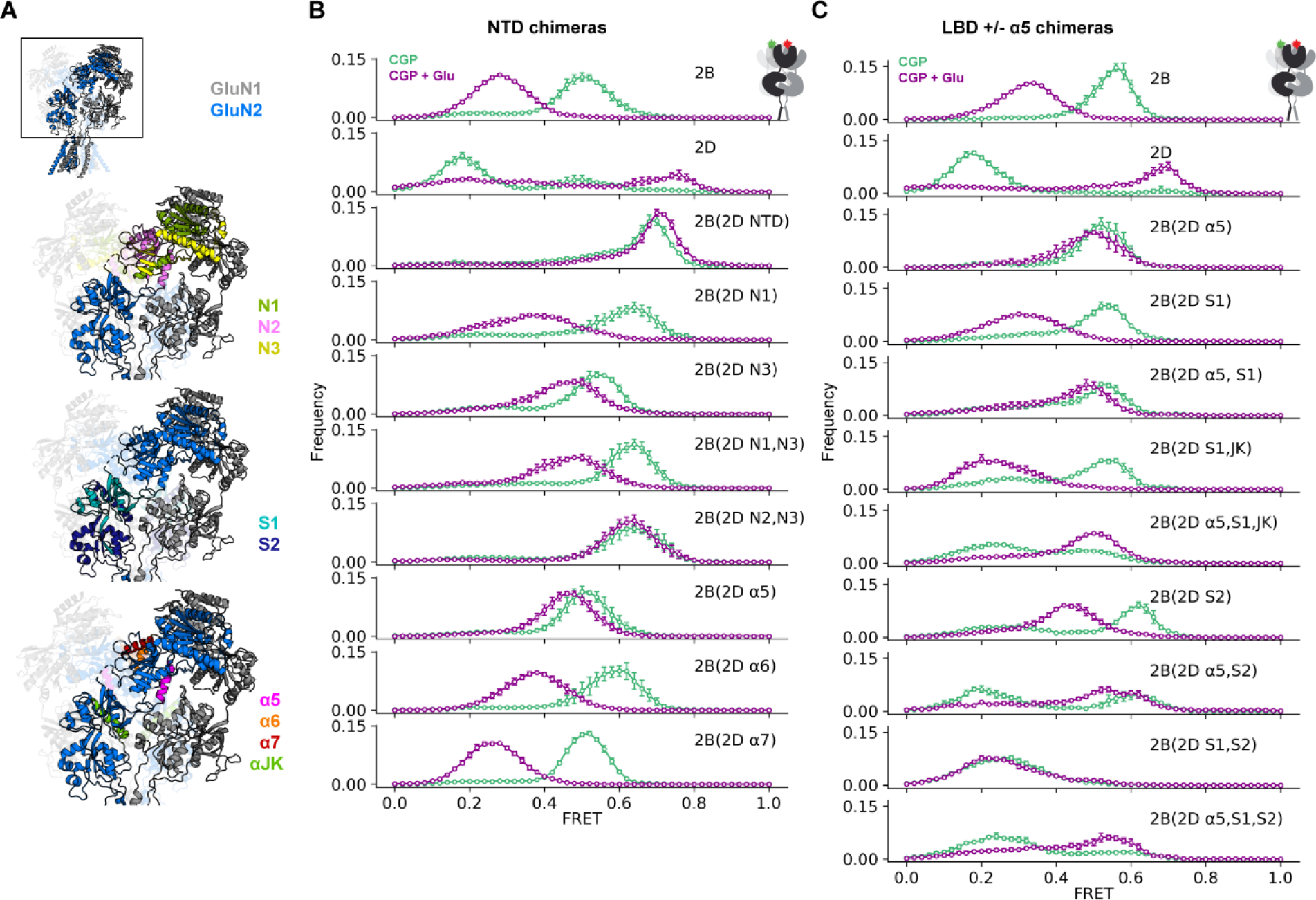
GluN2B/2D chimeras reveal structural determinants of subtype-specific conformational dynamics. (A) GluN1a/GluN2B glycine/glutamate structure (PDB 7SAA^72^) showing the regions swapped in chimeric GluN2 receptors. (B-C) Histograms of smFRET distributions indicating mean and S.E.M. for ligand-dependent conformations determined from inter-subunit FRET between GluN1(W56TAG) NTD paired with chimeric GluN2 subunits (according to scheme in Fig. S2), which transplant pieces of GluN2D into GluN2B, focusing on the NTD (B) and LBD (C) in 3 μM CGP78608 and 1 mM glutamate (purple) and apo-like (zero added glycine, in 3 μM CGP78608; green) conditions. Labeling sites indicated with green and red stars on cartoon insets at top right of (B) and (C). Total receptor number for each condition listed in Table S1. Donor (Alexa Fluor 555) and acceptor (Alexa Fluor 647) dyes imaged at 10 fps. See also Figure S5.

As the GluN2D NTD did not transfer the characteristic GluN2D splayed apo-like resting state, we examined the LBD, still using the GluN1(W56TAG) FRET sensor to monitor inter-NTD proximity. Substitution of the GluN2D LBD into GluN2B [2B(2D S1, S2)] did, in fact, result in a lower FRET, splayed, resting state in CGP, which resembled that of GluN2D (**Fig. 3C**). However, unlike the NTD swap, it did not transfer GluN2D-type NTD convergence in glutamate. This suggested that, compared to their respective GluN2B counterparts, the GluN2D LBD favors the splayed resting state and the α5 helix of the GluN2D NTD favors compact conformations when glutamate is bound. Indeed, substitution of both the GluN2D LBD and α5 helix [2B(2D α5, S1, S2)] results in both a splayed resting state and convergence with glutamate (**Fig. 3C**). Subdivision of the LBD showed that, whereas the GluN2D S1 along with helices J and K of S2 [2B(2D S1, JK)] have limited effect on their own, they are sufficient to impart the glutamate-induced NTD convergence of the entire GluN2 LBD when combined with the α5 helix of the NTD [2B(2D α5, S1, JK)] (**Fig. 3C**). This chimeric analysis indicates the critical nature of the α5 helix in conformational rearrangements of GluN2B receptors.

The GluN2B, NTD α5 helix interacts with the GluN1 LBD loop 2 in an apparently activation-state dependent manner.^53^ We tested the possibility that this interaction plays a part in the stabilization of the low FRET glutamate-alone conformation of the GluN1 NTDs (NTD-separated) in the GluN2B receptor. We monitored inter-GluN1(W56TAG,R489-K496GG) NTD proximity in receptors with GluN2B or GluN2D subunits and GluN1 subunits, where GluN1 LBD loop 2 residues 489-496 were replaced by a short flexible linker consisting of two glycine residues (**Fig. 4A, red loop**), which should eliminate interaction between GluN1 LBD loop 2 and GluN2 NTD α5. In the GluN2B receptor, the characteristic NTD separation with glutamate is greatly reduced by this loop 2 alteration (**Fig. 4B**), similar to what we observed with substitution of the GluN2D α5 (**Fig. 3**). This suggests that α5-loop 2 interaction stabilizes the NTD-separated conformation in glutamate bound GluN2B receptors. In contrast, in the GluN2D receptor, this perturbation has only a minor effect on the glutamate state observed at the NTD (**Fig. 4C**). In neither case is there an effect on the resting conformation (**Fig. 4B, C**). These observations suggest that interaction between GluN1 loop 2 and GluN2B α5 couples the GluN2B NTD to the GluN1 LBD, providing a mechanism through which glutamate binding to the GluN2 subunit allosterically regulates the conformation of the GluN1 subunit.

**Figure 4.**
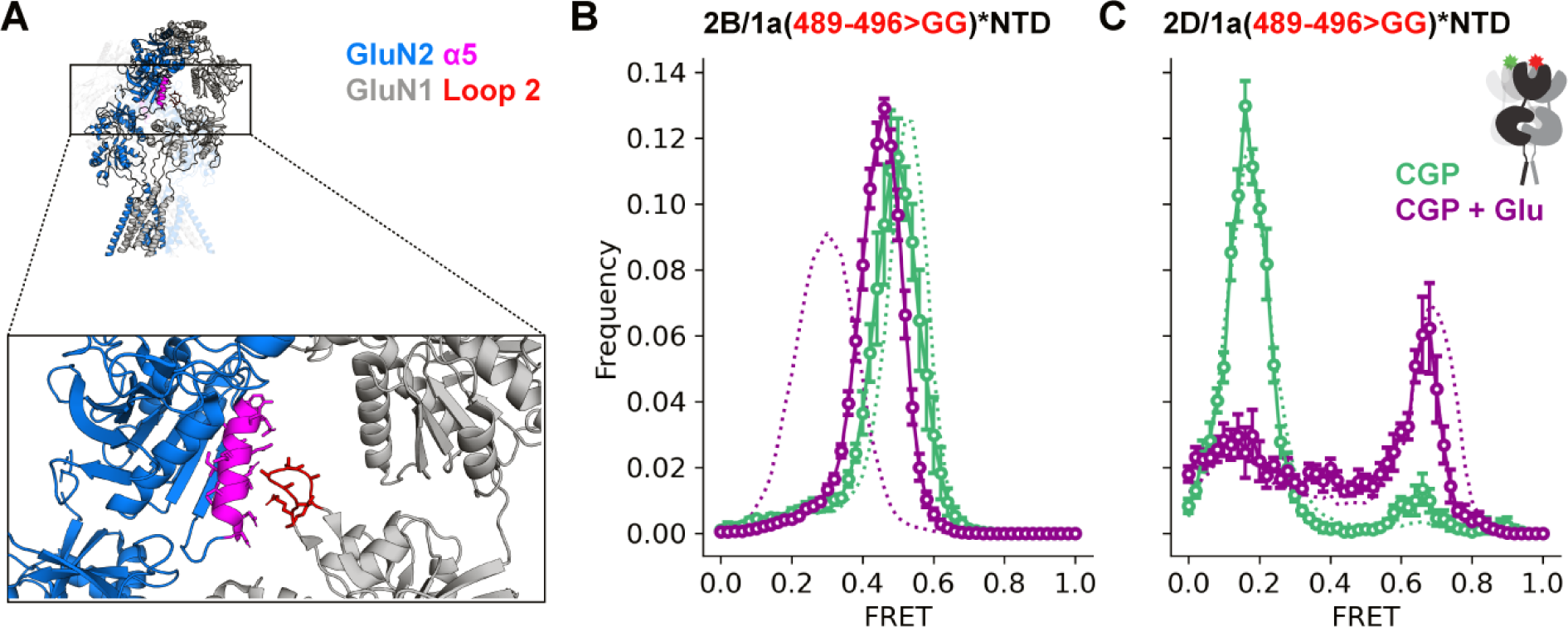
GluN1 loop 2 deletion affects glutamate-bound NTD conformation in GluN2B but not GluN2D receptors. (A) GluN2 α5 helix (magenta) and residues 489-496 of GluN1 Loop 2 (red) in the GluN1a/GluN2B glycine/glutamate structure (PDB 7SAA^72^) (B-C) Histograms of smFRET distributions indicating the mean and S.E.M. for ligand-dependent conformations determined from inter-subunit FRET between GluN1(W56TAG,R489-K496GG) NTD paired with GluN2B (B) or GluN2D (C) in 3 μM CGP78608 and 1 mM glutamate (purple) and apo-like (zero added glycine, in 3 μM CGP78608) conditions. Labeling sites indicated with green and red stars on cartoons. Total receptor number for each condition listed in Table S1. Donor (Alexa Fluor 555) and acceptor (Alexa Fluor 647) dyes imaged at 10 fps.

## Discussion

We show that GluN2 subunits differentially determine the conformational trajectory from resting to agonist-bound states in diheterotetrameric NMDA receptors. Using smFRET allowed us to interrogate fluctuation between states in the activation pathway that cannot be detected directly with electrophysiology and that have thus far not been detected with structural approaches. Our results provide insight into the conformational basis of subtype-specific allosteric control of NMDA receptor gating and help explain why receptors with higher P_o_ (GluN2A and GluN2B) exhibit lower agonist potency and show more desensitization than those with lower P_o_ (GluN2C and GluN2D).

We find that, along the activation pathway, in absence of agonist, or in either glutamate or glycine alone, low P_o_ GluN2C and GluN2D receptors favor splayed NTD conformations, with dynamic GluN1 LBDs that broadly sample diverse orientations. In contrast, higher P_o_ GluN2A and GluN2B receptors, which favor compact conformations, with more stable GluN1 LBD conformation and compact NTDs. In addition, in GluN2A and GluN2B receptors, glutamate promotes occupancy of more splayed conformations and glycine of more compact conformations, but in GluN2C and GluN2D receptors both agonists promote occupancy of more compact conformations. However, in the low P_o_ GluN2D, the fully agonist-bound receptor exhibits a super-compact NTD conformation, also observed recently in structural studies.^38,39^

Our results suggest that distinct occupancy of three common classes of conformations (splayed, compact, and super-compact) in apo, partially, and fully liganded conformations in different receptor subtypes gives rise to their distinct properties (**Fig. 5A**). The splayed and super-compact conformations in the low P_o_ GluN2D compared to the compact conformations of the high P_o_ GluN2A, suggest that splayed and super-compact conformations are nonproductive low P_o_ conformations and compact conformations include pre-open and open states. The association of splaying in GluN2C and GluN2D receptors with nonproductive states is consistent with earlier findings that associate splayed conformations with inhibited states including the reduction of macroscopic current following MTSET modification to prevent transition from splayed to compact conformations in the GluN2C receptor.^38^ Structural studies have also demonstrated that negative allosteric modulation by Zn^2+^ and H^+^ in the GluN2A receptor and orthosteric antagonism with D- 2-Amino-5-phosphonovaleric acid and dichlorokynurenic acid in the GluN2B receptor lead to occupancy of conformations with ruptured upper-LBD interfaces, swapped LBD arrangement, and splayed NTDs.^34,50^ In addition to the entry of unliganded or partially liganded GluN2D receptors into splayed non-productive states, from which it may take some time to emerge, the uniquely super-compact conformation of its fully liganded state could further reduce P_o_ in light of the earlier demonstration that cross-linking of the D1-D1 intersubunit interface within GluN1/GluN2A LBD dimers, which will increase receptor compaction, drastically reduces P_o._^54^

**Figure 5.**
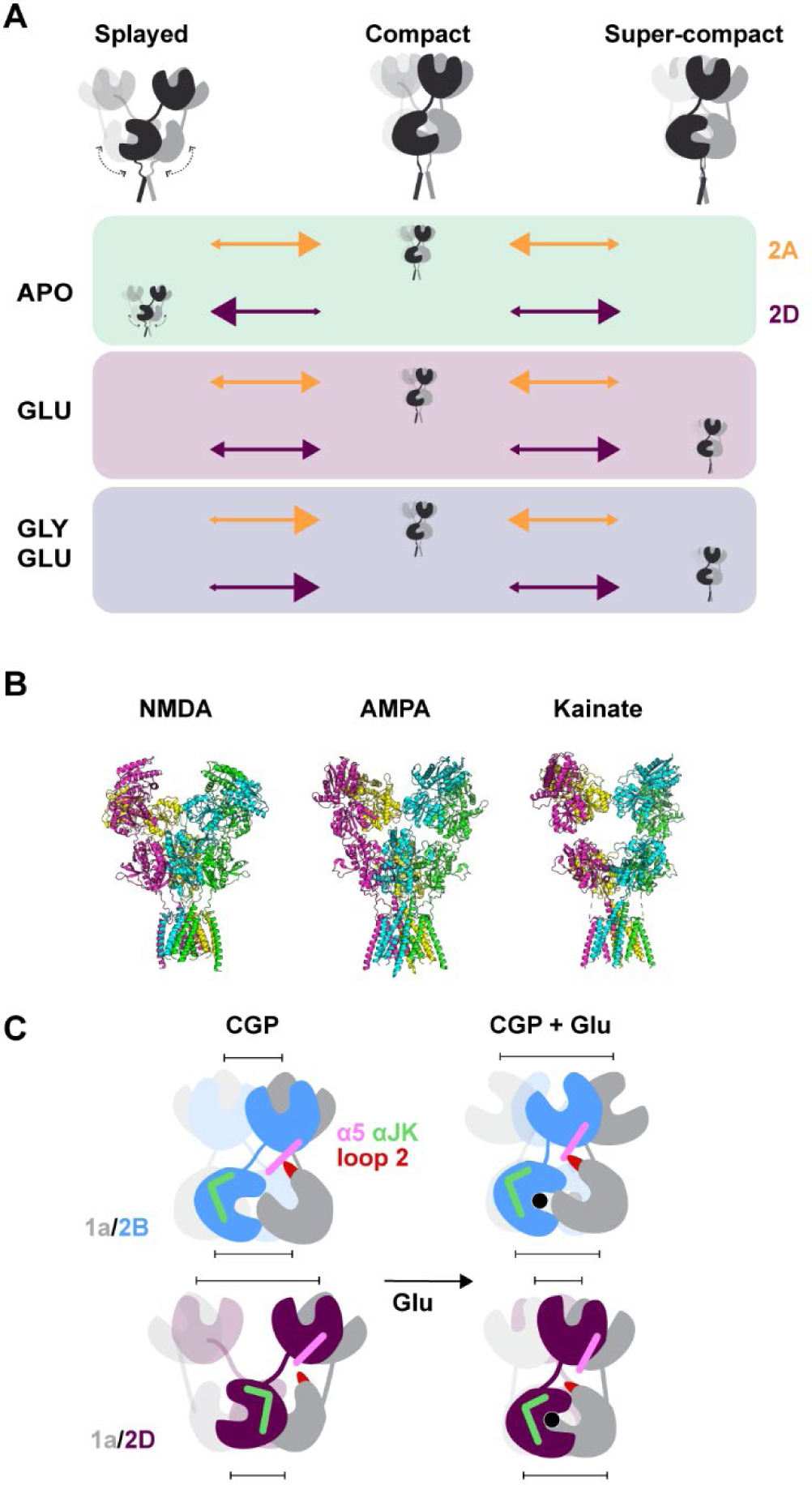
Model of conformational basis of GluN2 dependent NMDAR gating. (A) Cartoons of splayed, compact, and super-compact conformations and orange (GluN2A receptors) and purple (GluN2D receptors) arrows indicating relative occupancy (with size of arrowhead) of each conformation in apo, glutamate-bound, and glutamate and glycine-bound receptors. Cartoon in each row indicates primary conformation occupied in each ligand/receptor condition. (B) Structures of ionotropic glutamate receptors (NMDAR, PDB 7EOS: GluN2A/GluN1 with glycine and glutamate;^37^ AMPAR: GluA2 with glutamate (and TARP gamma-5 not shown) PDB 7RZ6;^73^ Kainate: GluK2/K5 with L-Glu, PDB 7KS3^74^) demonstrating increased NTD-LBD interaction in NMDA receptors. (C) Representation of key structural regions identified in chimeric analysis and their involvement in differences between GluN2B and GluN2D conformations in apo-like and glutamate-bound conformations. In GluN2B receptors interactions between the α5 helix of the GluN2 NTD and loop 2 of the GluN1 LBD result in occupancy of splayed conformations when glutamate is bound. In GluN2D receptors this interaction does not play a key role in observed conformations. Differences in helices J and K in the GluN2 LBD contribute to the stability of compact conformations.

As GluN2A (and GluN2B) receptors occupy primarily compact conformations in their resting state, they are primed to activate quickly upon agonist-binding. Synchronous openings will result in a large initial macroscopic current, which will decrease to a steady-state level that reflects the balance between bursts of openings and desensitized states that produce long-lived inter-burst closures, yielding macroscopic desensitization. In contrast, in their resting and partially liganded states, GluN2C and GluN2D receptors favor splayed conformations with mobile LBDs, which likely do not exert force efficiently on the gate; agonist binding likely generates delayed and therefore asynchronous openings, resulting in a slow-rising current of small-magnitude, directly to steady-state, with no evident macroscopic desensitization. Our interpretation of our results is that this steady-state current is particularly small in the GluN2D receptor because super- compaction in the fully liganded state results in a very low P_o_.

In AMPA and kainate receptors, which have little contact between the LBD and NTD layers (**Fig. 5B**), rupture of the upper LBD interface results in release of tension between the agonist-bound closed LBD and the channel gate in the TMD, resulting in desensitization.^2,55,56^ NTD splaying and LBD interface rupture in NMDA receptors also ostensibly represents the transition to nonproductive or desensitized states; however these conformations are less common in fully- agonist bound receptors, potentially owing to the extensive NTD-LBD contacts in all four GluN2 subtypes (**Fig. 5B**). Increased occupancy of more splayed conformations in GluN2A and GluN2B in partially-liganded glutamate bound receptors compared to the resting and glycine-bound conformations also provides an explanation for negative-cooperativity between glutamate and glycine binding, which gives rise to low agonist potency and glycine-dependent desensitization.^51,57–59,52,43^ In contrast, GluN2C and GluN2D receptors, which have higher agonist potency, glycine and glutamate both promote occupancy of more compact conformations. We find it interesting to consider the relationship between the two types of nonproductive conformations that we observe and the two desensitized states proposed by kinetic models.^60,61^ Increased occupancy of the super-compact conformation in GluN2A by disulfide crosslinking of the dimer interface in GluN1/GluN2A abolishes the less prevalent of the two desensitized states,^54^ suggesting it could correspond to the splayed conformation which GluN2A receptors only occupy appreciably in the absence of glycine in our assay.

Finally, our chimeric analysis suggests that the super-compaction of the GluN2D receptor is caused by the GluN2D NTD and that interaction between GluN2 NTD α5 and GluN1 LBD loop 2 receptor regulates glutamate-induced rearrangements in GluN2B. Previous studies indicate that, during activation, GluN1 LBD rolls forward and that its loop 2 moves down towards the GluN2B LBD.^33,36^ Indeed, crosslinking GluN1 loop 2 ‘higher up’ on the GluN2B α5 silences receptors but crosslinking ‘lower down’ increases P_o_, suggesting that activation rolling of the GluN1 LBD slides its loop 2 ‘down’ the GluN2B NTD α5 helix.^53^ Our chimeric analysis suggests that this rolling of the GluN1 LBD may be inhibited by glutamate binding through GluN1 loop 2 - GluN2 α5 interaction in GluN2B but not in GluN2D receptors, which demonstrate less potential for interaction between α5 and loop 2 (**Figs. 5C, S5D,E**). However, it remains possible that this interaction also plays a role in GluN2D receptors, which is not apparent with our NTD sensor due to differences in inter- GluN2 and GluN2-GluN1 interactions at the level of the NTD.

We additionally find a role for GluN2 S2 helices J and K in the upper lobe of the GluN2 LBD in regulating occupancy of splayed conformations. In compact conformations in both GluN2B and GluN2D, these helices are positioned to form inter-subunit interactions with the upper and lower lobes of alternate adjacent GluN1 subunits as well as intra-subunit interactions with S1 in the upper lobe of the GluN2 subunit **(Fig. S4B,C**). In contrast, in splayed conformations of antagonist-bound GluN2B receptors, the interfaces with adjacent GluN1 subunits are no longer present **(Fig. S4A**). If this also occurs in glutamate-bound splayed receptors, it may favor closed-cleft LBD conformations and contribute to higher glutamate affinity in splayed conformations as seen in desensitized AMPA receptors.^62–64^ Sequence variation in these helices in different GluN2 subunits likely affects the stability of interactions at each of these interfaces. The role we identify for these helices in subtype-specific regulation of receptor compaction is also of interest given their close proximity to both the hinge-loop region involved in deactivation time course^17^ and to the peripheral M4 transmembrane segment, which is known to play a subunit-specific regulatory role in gating and desensitization.^65,66^

Our work provides a conformational explanation for the distinct physiological properties observed in NMDA receptor subtypes. Our findings also suggest that factors that bias occupancy between splayed, compact and super-compact conformations could transition NMDA receptors between high and low P_o_ modes. It is intriguing to consider that endogenous modulation by binding partners such as auto-antibodies,^67^ other receptors and adhesion proteins found at synapses^68–71^ may modulate function by differentially favoring different states of compaction and that drug design that takes this into account could provide subtype and agonist-conformation specific modulators.

## Acknowledgments

We thank Howard Hang for providing the Mm-PylRS-AF/Pyl-tRNACUA plasmid, Naomi Latorraca for assistance with PyMOL, Naomi, Yawei Yu and other members of the Isacoff lab for critical discussions of data, Shimon Yudovich for assistance with epifluorescence imaging, and Lilia Garcia for generation of lab stocks used for molecular biology. This work was supported by the National Institutes of Health (R01NS119826 and R01 GM117051) and Nan Fung (Pivotal) Life Sciences (#5584649 fund R47470000) to E.Y.I. E.Y.I. is a Weill Neurohub Investigator.

## Author contributions

J.B. and C.S. performed molecular biology to generate modifications of GluN1 and GluN2 plasmids; C.H. performed molecular biology and incorporation tests to improve unnatural amino acid incorporation efficiency; V.V. provided insight from NMDAR smFRET experience including suggestion of W56TAG incorporation site; J.B. performed cell culture and transfections; P.R.F.M. performed patch-clamp recordings; J.B. conceptualized and performed smFRET imaging experiments and analysis with training and guidance from C.H.; E.Y.I. acquired funding, supervised the project and co-wrote the manuscript with J.B. with input from all authors.

## Declaration of Interests

The authors declare no competing interests.

## RESOURCE AVAILABILITY

### Lead contact

Requests for further information, resources, or reagents should be directed to and will be completed by the lead contact, Ehud Isacoff.

### Materials availability

Plasmids generated in this study will be made available upon request.

### Data and code availability

All data reported in this paper will be shared by the lead contact upon request. This paper does not report significant original code.

Any additional information required to reanalyze the data reported in this paper is available from the lead contact upon request.

## METHODS

### DNA constructs and site-directed mutagenesis

Amino acids and sites of mutations are numbered according to the wild type full length *rattus norvegicus* proteins (accession codes: BAA02498.1 (GluN2A), NP_036706.1 (GluN2B), XP_006247771.1 (GluN2C), NP_073634.2 (GluN2D)) beginning with methionine as 1. For GluN2 constructs, a flexible linker followed by a Human influenza hemagglutinin (HA) tag (GGGGS- YPYDVPDYA) was inserted immediately prior to the stop codon in the full-length protein. Chimeric GluN2 subunits (according to scheme in Fig. S2 created using Boxshade^75^ with T-Coffee alignment^76^) involving large exchanged regions were generated using Gibson assembly and smaller regions and other modifications of GluN1 and GluN2 subunits were generated using PCR mutagenesis. Modifications in the Mm-PylRS-AF/Pyl-tRNACUA plasmid were generated using gBlocks (IDT) and Gibson assembly and included insertion of three additional copies of the pyrrolysyl-tRNA as well as insertion (the translation elongation factor EF1A and a nuclear export sequence MACPVPLQLPPLERLTLD from the HIV-1 transactivating protein Rev) and removal (FLAG) of elements upstream and insertion of the woodchuck hepatitis virus post-transcriptional regulatory element downstream of the Pyrrolysyl-tRNA Synthetase(AF).

### Cell culture and transfection

Human embryonic kidney 293T (ATCC: CRL-3216) were cultured in Opti-MEM (Gibco 31985070) supplemented with 5% fetal bovine serum on 3 µg/mL collagen coated plates at 37°C in 5% CO_2_. For smFRET experiments cells were cultured for approximately 3-25 passages before cells were seeded in 9.6cm^2^ 6-well plates coated with 0.5 mg/mL poly-L-lysine. At ∼80% confluency, up to 6 wells of each construct combination were transfected. Media in each well was first replaced with 1 mL of Opti-MEM transfection media supplemented with 3% FBS, 20 mM MgCl2, 50 µM 5,7- dichlorokynurenic acid (5,7-DCKA), 800 µM D,L-2-amino-5-phosphonopentanoic acid (D,L-APV) and 20 µM ifenprodil. After 20 minutes, a mixture of 250 µL Opti-MEM, 5 µL lipofectamine and 5 µg of DNA (at a ratio of 5:1:5 of GluN1-1a plasmid with TAG stop codon at position W56 or D677; GluN2 subunit with a C-terminal GGGGSS linker followed by an HA tag (YPYDVPDYA); 4XpylT- EF1a-NES-Mm-PylRS(AF)-WPRE amber suppression plasmid). 12.5 µL of 25 mM Trans-cyclooctene lysine (TCOK) (SiChem) was added to each well for a final concentration of ∼250 µM. TCOK stock was prepared at 100 mM in 0.2 M NaOH, 15% DMSO and was diluted 1:4 in 1 M HEPES before addition to cell media. Media was not changed again prior to receptor labeling on the day of imaging. For patch-clamp experiments the same procedure was followed with the following exceptions: cells were seeded on 18mm acid-washed borosilicate glass coverslips coated with poly-L-lysine (1 mg/mL) at a low density in 3.5cm^2^ 12-well plates; transfection occurred ∼5 hours later with transfection media consisting of 1.5% FBS, 20 mM MgCl2, 50 µM 5,7- DCKA and 400 µM D,L-APV. The mixture added after 20 minutes included 100 µL Opti-MEM, 2 µL lipofectamine and 1200 ng of DNA (500 ng GluN1-1a(W56TAG) or GluN1-1a(D677TAG), 100 ng GluN2 subunit with a C-terminal GGGGSS linker followed by an HA tag (YPYDVPDYA), 500 ng 4XpylT-EF1a-NES-Mm-PylRS(AF)-WPRE amber suppression plasmid, and 100 ng tdTomato).

### Patch-clamp electrophysiology

Patch-clamp recordings were performed 16-24 hours following transfection. Each coverslip was washed in extracellular buffer (pH 7.4 with NaOH) containing, in mM: 160 NaCl, 2.5 KCl, 10 HEPES, 0.2 EDTA, 0.7 CaCl_2_, 1 MgCl_2_ and labeled in 300 nM of pyrimidyl-tetrazine-Alexa647 (JENA Biosciences) for 15-20 minutes. Following labeling, coverslips were transferred onto a recording chamber mounted on an Olympus IX71 inverted microscope, with a Mg^2+^-free version of the extracellular buffer additionally containing 100 µM glycine. Cells expressing TdTomato were identified using a DG-4 light excitation system (Sutter instruments). Voltage-clamp recordings were obtained using borosilicate glass pipettes with 4-6MΩ resistance filled with the following intracellular solution (in mM): 120 gluconic acid, 15 CsCl, 10 BAPTA, 10 HEPES, 3 MgCl_2_, 1 CaCl_2_, and 2 ATP-Mg salt (pH-adjusted to 7.2 with CsOH). After establishing whole-cell configuration, cells were held at -70mV. Liquid junction potential was not corrected. A second borosilicate glass pipette (2-4 MΩ) was loaded with 1 mM glutamate and positioned directly in front of the patched cell. A gentle positive pressure was applied to locally perfuse glutamate. Data was acquired using a CV203BU head stage, Axopatch 200B amplifier (Molecular Devices), and a Digidata 1440 acquisition board controlled with pCLAMP software, with data sampled at 10Khz, Bessel filtered at 4Khz.

### UAA-mediated NMDAR labeling and solubilization

Receptor labeling was performed on cells 24-48 hrs following transfection. Transfection media was removed and each well was washed twice in 1 mL extracellular buffer (pH 7.4 with NaOH) containing, in mM: 160 NaCl, 2.5 KCl, 2 CaCl_2_, 1 MgCl_2_, and 10 HEPES. 450 µL labeling solution containing 300 nM of each tetrazine-Alexa555 and tetrazine-Alexa647 (JENA Biosciences) in extracellular buffer was added to each well. The 6-well plate was then placed in an opaque container containing 4°C ddH2O and rocked gently at room temperature for 20 minutes. Following labeling, each well was washed with 1 mL extracellular buffer, 1 mL PBS (-/- Ca2+/Mg2+), and 1 mL PBS + 1 mM PMSF (Thermo Scientific) was added. Cells were incubated at 4°C for ∼5 minutes before being gently collected from the bottom of the well with a cell-scraper. Cell suspensions were spun down at 5000g for 5 minutes to pellet cells. Each pellet was resuspended in a lysis buffer containing (150 mM NaCl, 20 mM Tris (pH 8.0), 1% lauryl maltose neopentyl glycol (LMNG), 0.1% cholesteryl hemisuccinate (CHS) (Anatrace), protease inhibitor cocktail (Thermo Fisher Scientific), 1 mM PMSF, 5% glycerol) and allowed to shake in the dark for 90 minutes at 4°C. Following lysis, lysate was spun at 16,000g for 20 minutes and supernatant containing detergent-solubilized receptors was collected. This supernatant was subjected to 3 additional spins in 50 kDa Amicon Ultra 0.5 mL buffer exchange columns using a modified imaging buffer (pH 8.0 w/ NaOH) consisting of (in mM) 160 NaCl, 2.5 KCl, 2 CaCl_2_, 10 MgCl_2_, 20 HEPES, and of 0.01% LMNG, 0.001% CHS to remove any remaining dye, inhibitors used in transfection, and glutamate.

### SiMPull receptor isolation and surface display

Imaging chambers for single-molecule experiments were prepared using aminosilane functionalized glass coverslips and slides. To prevent non-specific binding, slides were passivated with mPEG (Laysan Bio) and coverslips were passivated with mPEG and biotin PEG16. 5-8 holes were drilled on each edge of a coverslip sized area of the slide prior to cleaning and passivation. Slides and coverslips were stored at -20°C until the day of each experiment. Double-sided adhesive was used to attach coverslips to slides and create several channels which were sealed with quick drying epoxy (Devcon) through which solutions could be flowed. On the day of each experiment, channels were incubated with 20 µg/ml NeutrAvidin (Thermo Fisher Scientific) for 15 min, followed by 1/100 biotinylated anti-HA antibody (Abcam, ab26228) for at least 1 hr. T50 buffer (50 mM NaCl, 10 mM Tris, pH 7.4) was used to dilute NeutrAvidin, anti-HA antibody as well as to wash each out of the chamber. Cell lysate was diluted (1-10x) and incubated in the imaging chamber (1-30 min) to achieve sparse mobilization. Unbound lysate was washed out extensively using a modified imaging buffer (pH 8.0 w/ NaOH) consisting of (in mM) 160 NaCl, 2.5 KCl, 2 CaCl2, 10 MgCl2, 20 HEPES and of 0.01% LMNG, 0.001% CHS.

### smFRET measurements

Receptors were imaged for smFRET in imaging buffer (pH 8.0 w/ NaOH) consisting of (in mM) 160 NaCl, 2.5 KCl, 2 CaCl2, 10 MgCl2, 20 HEPES, 50 glucose, 0.01% LMNG, 0.001% CHS, 5 Trolox, and 2 protocatechuic acid. 50 nM protocatechuate-3,4-dioxygenase and any ligands were added into a total volume of 100 µL of imaging buffer immediately before it was loaded into imaging chamber. Micro-Manager 2.0.0-beta3^77^ was used to control excitation of donor fluorophores with a 532 nM laser (Cobolt) and acquisition with an objective-based TIRF microscope (1.65 NA, 60x Olympus) and Photometrics Prime 95B sCMOS camera at 100-ms frame rate. For each condition, at least 4 movies were collected in different regions of a single imaging channel. All experiments were repeated at least twice on separate days with similar results. Data included in individual figure panels was collected on the same day.

## QUANTIFICATION AND STATISTICAL ANALYSIS

### smFRET analysis

smFRET data was processed using SPARTAN,^78^ where traces were extracted from acquired movies using the GetTraces module, subjected to selection in AutoTraces (default criteria except FRET lifetime > 50) and subsequent manual selection to ensure single-step bleaching of each fluorophore, constant total fluorescence, and global anti-correlation between donor and acceptors. From SPARTAN, display histograms (50 frames, bin size 0.02) and traces were exported (ForOrigin) and imported into a Jupyter notebook^79^ in which histograms of traces in individual movies (occasionally traces from up to 3 movies were combined in cases where expression was low) were averaged. S.E.M. was also calculated across the sets of traces from individual movies and traces and histograms were plotted. Median and quartiles were calculated as the FRET value corresponding to the first bin with over 25, 50 and 75% of cumulative counts for individual movies and averaged for each condition. Trimodal gaussian fitting was achieved using scipy.optimize.curve_fit with gaussians each defined as A*np.exp(-(x- mu_n)**2/(2*sigma**2)) with initial A=0.05 and sigma=0.01 and fixed means (mu_1=0.3, mu_2=0.5, mu_3=.7). Area under each curve was calculated using simpson’s rule (scipy.integrate.simpson) and reported as a percentage of the area under the trimodal fit.

## Notes

### Competing Interest Statement

The authors have declared no competing interest.

