## Supplementary Data for "Conformational basis of subtype-specific allosteric control of NMDA receptor gating"

### Supplemental Figures

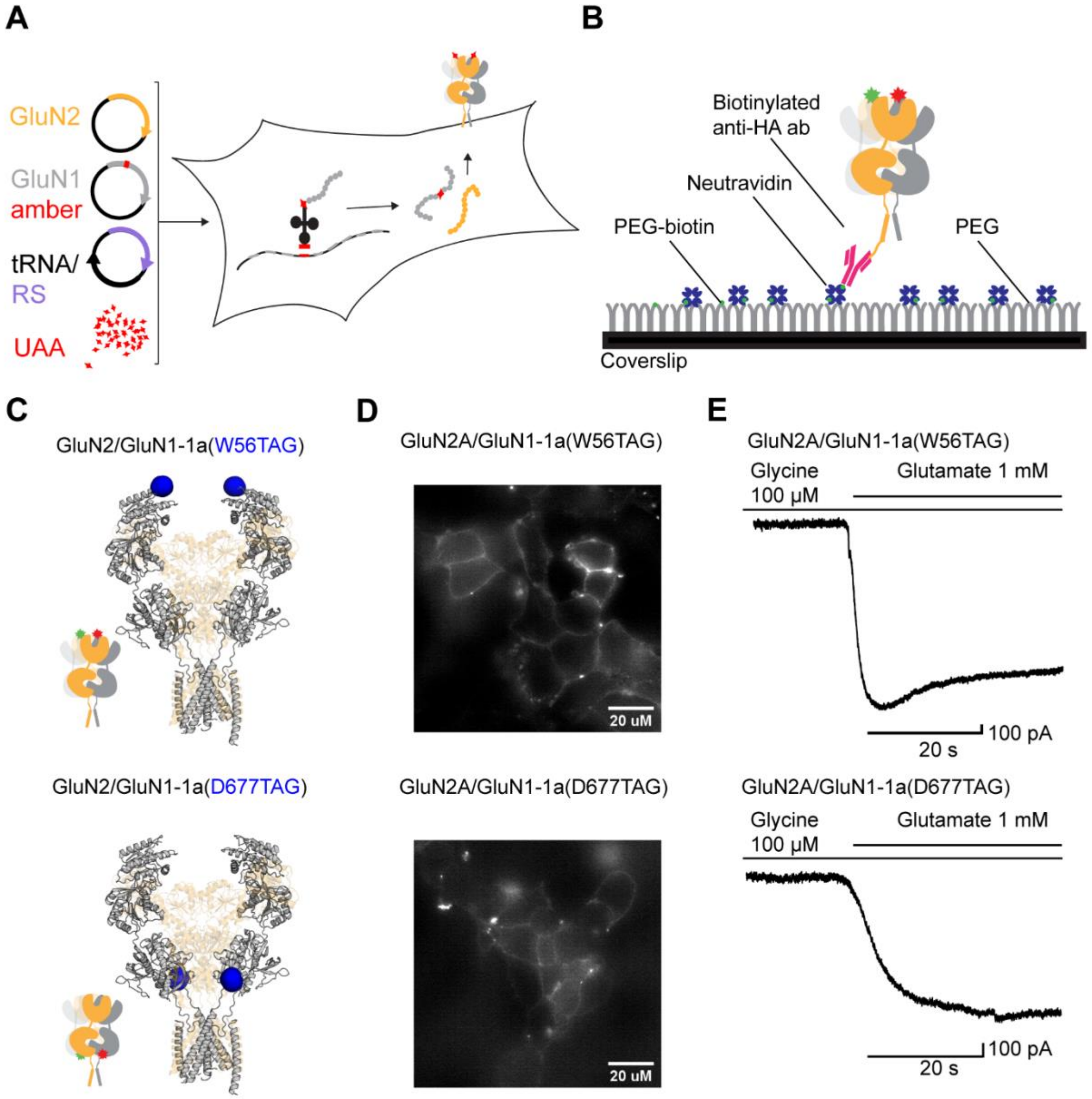


**Figure S1. Expression and labeling of TCOK-incorporated NMDA receptors** (A) Schematic of expression of NMDA receptors with site-specifically incorporated unnatural amino acids. Diagram inspired by Klippenstein et al. 2017[1] (B) Schematic of labeled immune-purified NMDA receptor in imaging chamber with GluN2 HA tag bound to biotinylated anti-HA antibody, which is bound to neutravidin which is bound to biotin-PEG (C) Structures (PDB 7EOS[2]) and cartoons depicting GluN1 labeling sites (D) Epifluorescence image of acceptor (tetrazine-AF647) labeled HEK293T cells expressing GluN1(W56TAG)/GluN2A (top) or GluN1(D677TAG)/GluN2A (bottom) receptors. (E) Representative whole cell patch clamp current traces in donor and acceptor labeled GluN1(W56TAG)/GluN2A (top) or GluN1(D677TAG)/GluN2A (bottom) receptors in HEK293T cells in response to pipette application of 1 mM glutamate in the presence of 100 µM glycine.

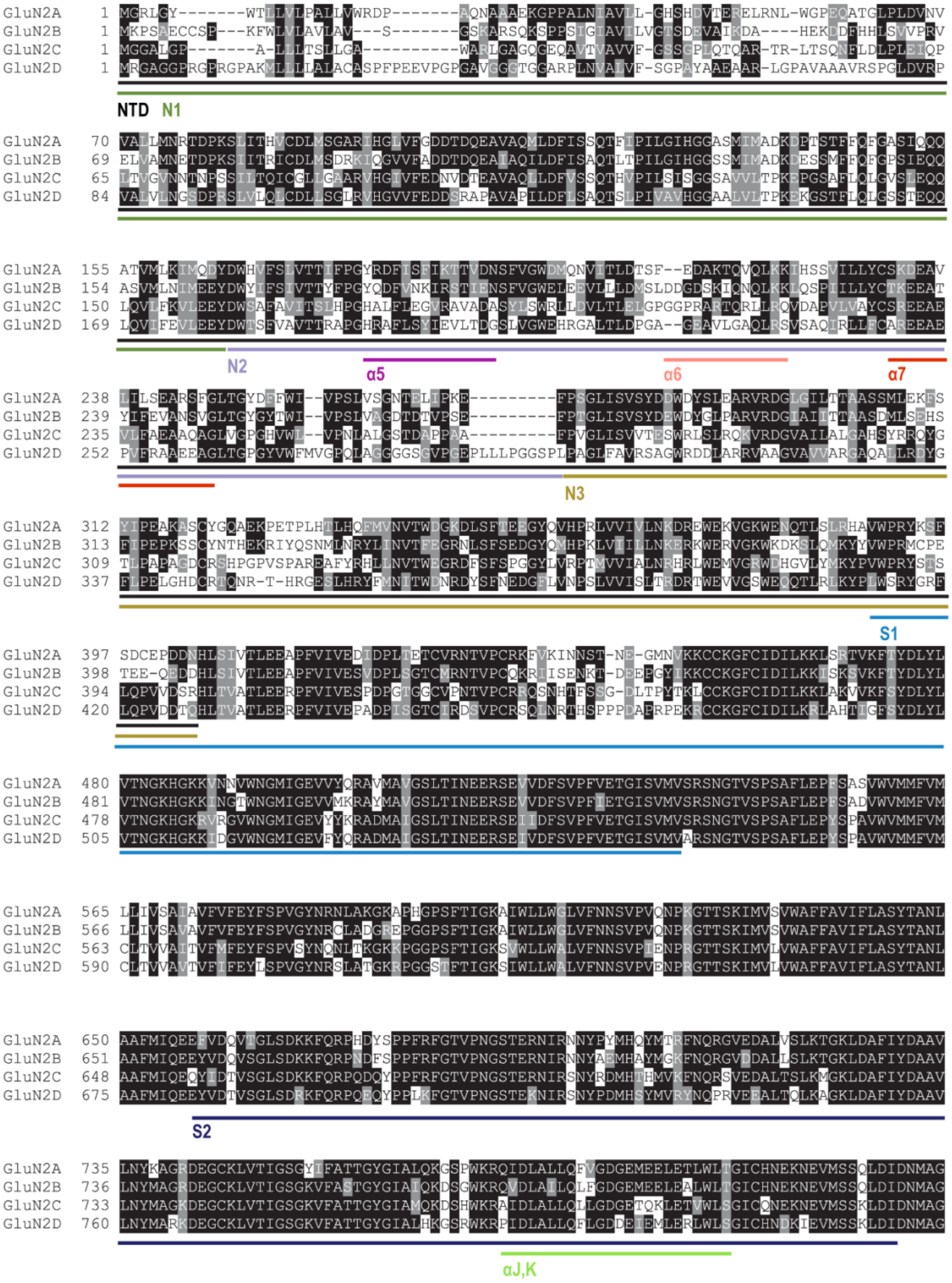


**Figure S2. GluN2 sequence alignment** Partial GluN2 sequence alignment (T-Coffee[3]) showing the regions that were exchanged in chimeric receptors, shaded with Boxshade.[4]

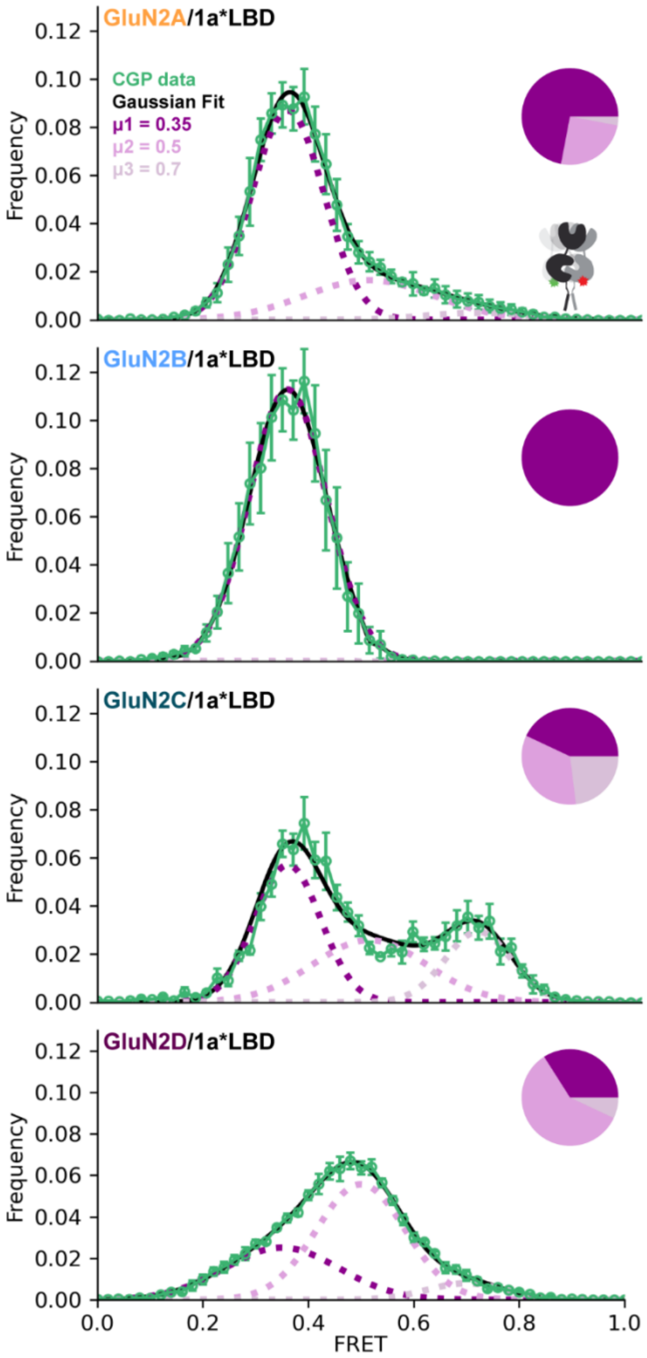


**Figure S3. Gaussian fits.** Trimodal Gaussian fits (black) of inter-subunit FRET distribution (green, mean +/- S.E.M., from Fig. 1A) between GluN1(D677TAG) LBDs in Apo-like state (zero added glycine, 3 μM GluN1 antagonist CGP78608, zero added glutamate) combined with each of the GluN2 subunits. Individual Gaussians centered at 0.35, 0.5, and 0.7 (dotted, shades of purple) with the corresponding percentages of area of the total fit in pie charts (right insets). Percentages, rounded to the nearest integer for Gaussians centered at FRET = 0.35: 72 (GluN2A), 100 (GluN2B), 43 (GluN2C), 34 (GluN2D); FRET = 0.5: 25 (GluN2A), 0 (GluN2B), 34 (GluN2C), 59 (GluN2D); FRET = 0.7: 3 (GluN2A), 0 (GluN2B), 23 (GluN2C), 7 (GluN2D).

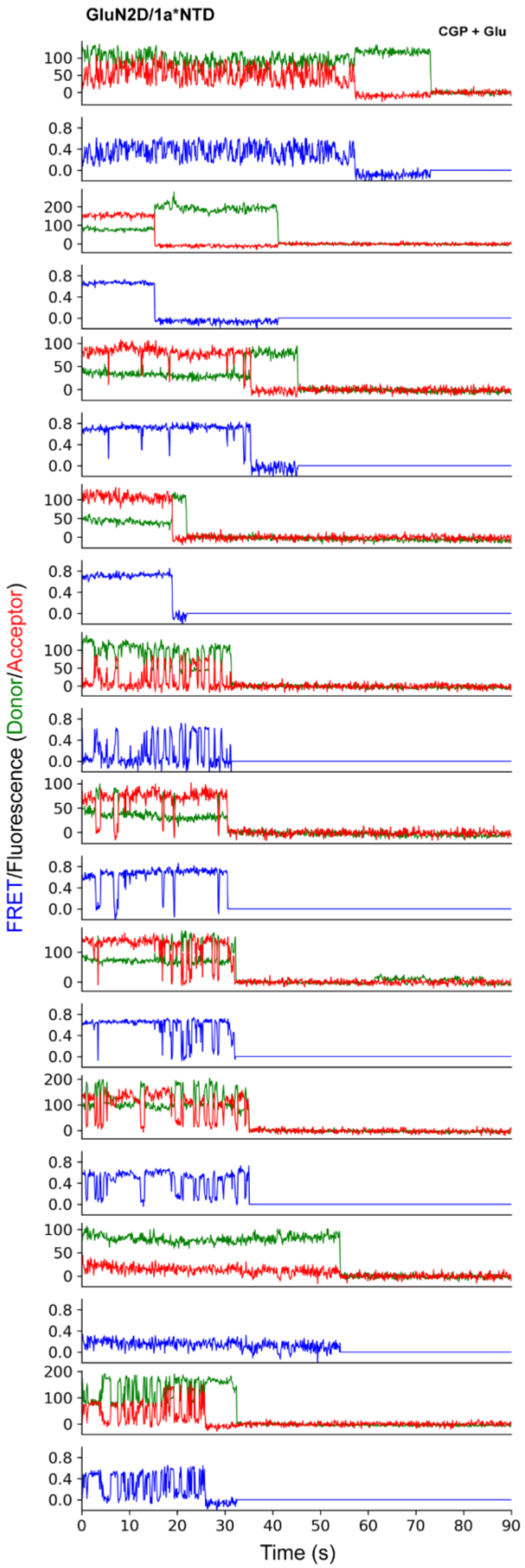


**Figure S4. Example traces showing interconversions between splayed, compact and super-compact states** Reporting inter-subunit FRET (blue) between GluN1(W56TAG) NTD paired with GluN2D in 3 µM CGP78608 and 1 mM glutamate. These traces are representative of and included in the CGP + Glu histogram in Figure 2H. Donor (Alexa Fluor 555; green) and acceptor (Alexa Fluor 647; red) dyes imaged at 10 fps.

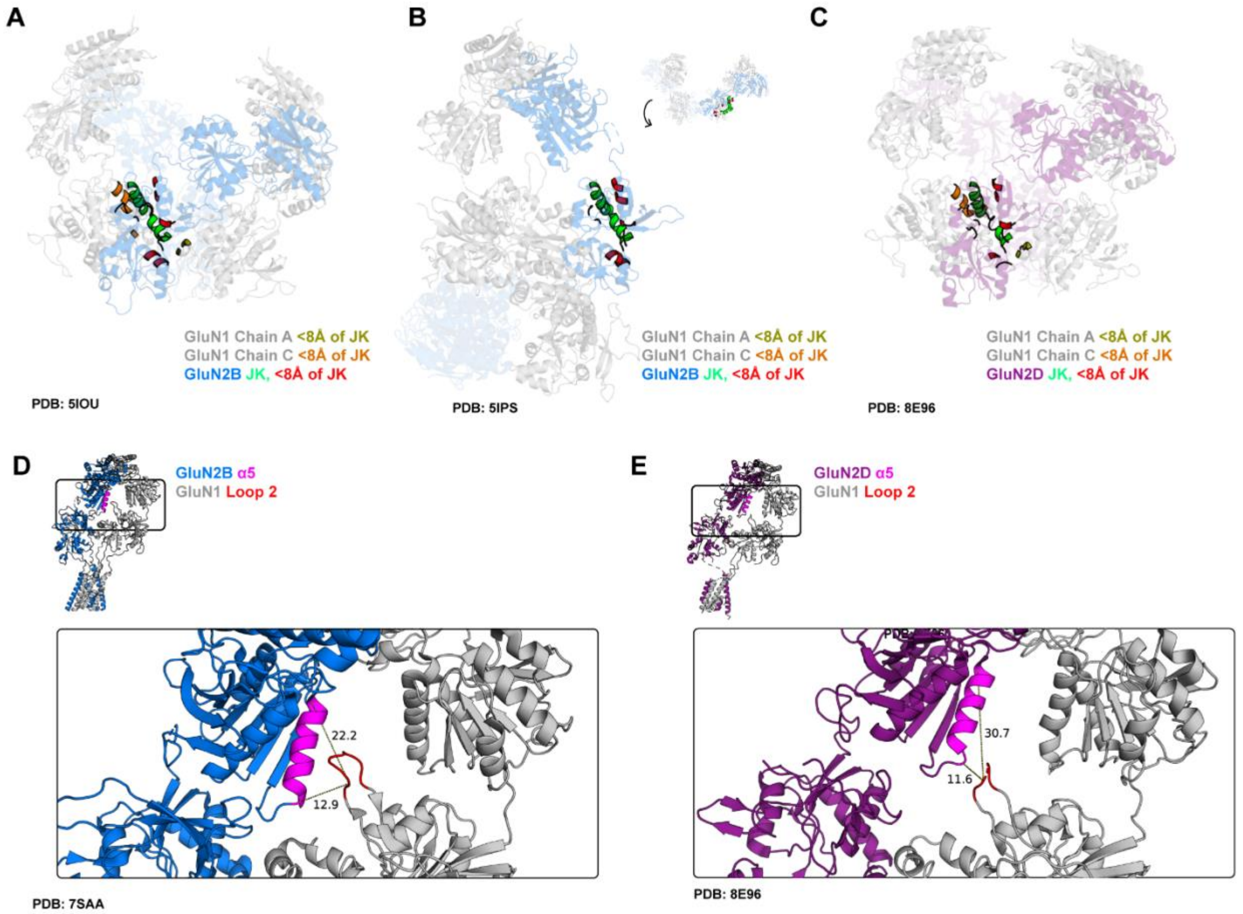


**Figure S5. Structures showing interfaces with regions identified in analysis of GluN2B/GluN2D receptor chimeras.** (A-C) Highlight of chimeric regions showing exchanged residues in helices J and K (green) and all residues within 8Å of backbone atoms of helices J and K, excluding those immediately adjacent in sequence (segments colored as indicated below structures) in GluN2B splayed (DCKA/D-APV, state 4) structure (PDB: 5IOU[5]) (A), GluN2B compact (glutamate/glycine) structure (PDB: 5IPS[5]) (B) and GluN2D super-compact (glutamate/glycine) structure (PDB: 8E96[7]) (C). (D,E) GluN2 α5 (magenta) – GluN1 loop 2 (red) interface in GluN2B (PDB: 7SAA[6]) (D) and GluN2D (PDB: 8E96[7]) (E). Whole structure above; blown up rectangular region below.

**Table S1. Total particle number included in histograms for each condition, by figure.**

| Figure | Construct | Condition | Total # Particles Included |
|---|---|---|---|
| 2 | 2B/1a(D677TAG) | 3uMCGP + 1mM Glu | 139 |
| 1,2,S3 | 2B/1a(D677TAG) | 3uM CGP | 101 |
| 1,2 | 2B/1a(D677TAG) | 100uM Gly + 1mM Glu | 226 |
| 2 | 2B/1a(D677TAG) | 100uM Gly | 243 |
| 2 | 2D/1a(D677TAG) | 3uMCGP + 1mM Glu | 172 |
| 1,2,S3 | 2D/1a(D677TAG) | 3uM CGP | 220 |
| 1,2 | 2D/1a(D677TAG) | 100uM Gly + 1mM Glu | 160 |
| 2 | 2D/1a(D677TAG) | 100uM Gly | 157 |
| 2 | 2A/1a(D677TAG) | 3uMCGP + 1mM Glu | 138 |
| 1,2,S3 | 2A/1a(D677TAG) | 3uM CGP | 157 |
| 1,2 | 2A/1a(D677TAG) | 100uM Gly + 1mM Glu | 116 |
| 2 | 2A/1a(D677TAG) | 100uM Gly | 246 |
| 2 | 2C/1a(D677TAG) | 3uMCGP + 1mM Glu | 88 |
| 1,2,S3 | 2C/1a(D677TAG) | 3uM CGP | 80 |
| 1,2 | 2C/1a(D677TAG) | 100uM Gly + 1mM Glu | 95 |
| 2 | 2C/1a(D677TAG) | 100uM Gly | 87 |
| 2 | 2B/1a(W56TAG) | 3uMCGP + 1mM Glu | 588 |
| 1,2 | 2B/1a(W56TAG) | 3uM CGP | 577 |
| 1,2 | 2B/1a(W56TAG) | 100uM Gly + 1mM Glu | 783 |
| 2 | 2B/1a(W56TAG) | 100uM Gly | 849 |
| 2,S4 | 2D/1a(W56TAG) | 3uMCGP + 1mM Glu | 266 |
| 1,2 | 2D/1a(W56TAG) | 3uM CGP | 116 |
| 1,2 | 2D/1a(W56TAG) | 100uM Gly + 1mM Glu | 290 |
| 2 | 2D/1a(W56TAG) | 100uM Gly | 229 |
| 2 | 2A/1a(W56TAG) | 3uMCGP + 1mM Glu | 425 |
| 1,2 | 2A/1a(W56TAG) | 3uM CGP | 452 |
| 1,2 | 2A/1a(W56TAG) | 100uM Gly + 1mM Glu | 330 |
| 2 | 2A/1a(W56TAG) | 100uM Gly | 416 |
| 2 | 2C/1a(W56TAG) | 3uMCGP + 1mM Glu | 236 |
| 1,2 | 2C/1a(W56TAG) | 3uM CGP | 276 |
| 1,2 | 2C/1a(W56TAG) | 100uM Gly + 1mM Glu | 355 |
| 2 | 2C/1a(W56TAG) | 100uM Gly | 335 |
| 3 | 2B(2D N1)/1a(W56TAG) | 3uMCGP + 1mM Glu | 206 |
| 3 | 2B(2D N1)/1a(W56TAG) | 3uM CGP | 215 |
| 3 | 2B(2D N3)/1a(W56TAG) | 3uMCGP + 1mM Glu | 608 |
| 3 | 2B(2D N3)/1a(W56TAG) | 3uM CGP | 487 |
| 3 | 2B(2D N1 N3)/1a(W56TAG) | 3uMCGP + 1mM Glu | 440 |
| 3 | 2B(2D N1 N3)/1a(W56TAG) | 3uM CGP | 186 |
| 3 | 2B(2D N2 N3)/1a(W56TAG) | 3uMCGP + 1mM Glu | 312 |
| 3 | 2B(2D N2 N3)/1a(W56TAG) | 3uM CGP | 373 |
| 3 | 2B/1a(W56TAG) | 3uMCGP + 1mM Glu | 812 |
| 3 | 2B/1a(W56TAG) | 3uM CGP | 566 |
| 3 | 2D/1a(W56TAG) | 3uMCGP + 1mM Glu | 322 |
| 3 | 2D/1a(W56TAG) | 3uM CGP | 184 |
| 3 | 2B(2D alpha5)/1a(W56TAG) | 3uMCGP + 1mM Glu | 641 |
| 3 | 2B(2D alpha5)/1a(W56TAG) | 3uM CGP | 487 |
| 3 | 2B(2D NTD)/1a(W56TAG) | 3uMCGP + 1mM Glu | 190 |
| 3 | 2B(2D NTD)/1a(W56TAG) | 3uM CGP | 182 |
| 3 | 2B(2D alpha6)/1a(W56TAG) | 3uMCGP + 1mM Glu | 671 |
| 3 | 2B(2D alpha6)/1a(W56TAG) | 3uM CGP | 548 |

| | | | |
|---|---|---|---|
| 3 | 2B(2D alpha7)/1a(W56TAG) | 3uMCGP + 1mM Glu | 592 |
| 3 | 2B(2D alpha7)/1a(W56TAG) | 3uM CGP | 617 |
| 3 | 2B/1a(W56TAG) | 3uMCGP + 1mM Glu | 317 |
| 3 | 2B/1a(W56TAG) | 3uM CGP | 270 |
| 3 | 2D/1a(W56TAG) | 3uMCGP + 1mM Glu | 169 |
| 3 | 2D/1a(W56TAG) | 3uM CGP | 145 |
| 3 | 2B(2D S2)/1a(W56TAG) | 3uMCGP + 1mM Glu | 192 |
| 3 | 2B(2D S2)/1a(W56TAG) | 3uM CGP | 156 |
| 3 | 2B(2D S1)/1a(W56TAG) | 3uMCGP + 1mM Glu | 408 |
| 3 | 2B(2D S1)/1a(W56TAG) | 3uM CGP | 299 |
| 3 | 2B(2D alpha5)/1a(W56TAG) | 3uMCGP + 1mM Glu | 124 |
| 3 | 2B(2D alpha5)/1a(W56TAG) | 3uM CGP | 150 |
| 3 | 2B(2D S1 S2)/1a(W56TAG) | 3uMCGP + 1mM Glu | 207 |
| 3 | 2B(2D S1 S2)/1a(W56TAG) | 3uM CGP | 270 |
| 3 | 2B(2D alpha 5 S1 S2)/1a(W56TAG) | 3uMCGP + 1mM Glu | 181 |
| 3 | 2B(2D alpha 5 S1 S2)/1a(W56TAG) | 3uM CGP | 210 |
| 3 | 2B(2Dalpha5 S1 JK)/1a(W56TAG) | 3uMCGP + 1mM Glu | 351 |
| 3 | 2B(2Dalpha5 S1 JK)/1a(W56TAG) | 3uM CGP | 342 |
| 3 | 2B(2Dalpha5 S1)/1a(W56TAG) | 3uMCGP + 1mM Glu | 314 |
| 3 | 2B(2Dalpha5 S1)/1a(W56TAG) | 3uM CGP | 244 |
| 3 | 2B(2Dalpha5 S2)/1a(W56TAG) | 3uMCGP + 1mM Glu | 108 |
| 3 | 2B(2Dalpha5 S2)/1a(W56TAG) | 3uM CGP | 107 |
| 4 | 2B/1a(W56TAG,489-496GG) | 3uMCGP + 1mM Glu | 684 |
| 4 | 2B/1a(W56TAG,489-496GG) | 3uM CGP | 563 |
| 4 | 2B/1a(W56TAG) | 3uMCGP + 1mM Glu | 348 |
| 4 | 2B/1a(W56TAG) | 3uM CGP | 380 |
| 4 | 2D/1a(W56TAG) | 3uMCGP + 1mM Glu | 119 |
| 4 | 2D/1a(W56TAG) | 3uM CGP | 151 |
| 4 | 2D/1a(W56TAG,489-496GG) | 3uMCGP + 1mM Glu | 107 |
| 4 | 2D/1a(W56TAG,489-496GG) | 3uM CGP | 97 |
